# STRT-seq-2i: dual-index 5′ single cell and nucleus RNA-seq on an addressable microwell array

**DOI:** 10.1101/126268

**Authors:** Hannah Hochgerner, Peter Lönnerberg, Rebecca Hodge, Jaromir Mikes, Abeer Heskol, Hermann Hubschle, Philip Lin, Simone Picelli, Gioele La Manno, Michael Ratz, Jude Dunne, Syed Husain, Ed Lein, Maithreyan Srinivasan, Amit Zeisel, Sten Linnarsson

**Author notes:** Corresponding authors. (A.Z.) (S.L.).

## Abstract

Single-cell RNA-seq has become routine for discovering cell types and revealing cellular diversity, but currently no high-throughput platform has been used successfully on archived human brain samples. We present STRT-seq-2i, an addressable 9600-microwell array platform, combining sampling by limiting dilution or FACS, with imaging and high throughput at competitive cost. We applied the platform to fresh single mouse cortical cells and to frozen post-mortem human cortical nuclei, matching the performance of a previous lower-throughput platform.

## Introduction

Single-cell RNA sequencing has become the method of choice for discovering cell types^1^,^2^ and lineages^3–5^, and for characterizing the heterogeneity of tumors^6^,^7^ and normal tissues such as lung^8^ and the nervous system^9^. Protocols with high levels of accuracy, sensitivity and throughput are now available commercially and from academia. Commonly used platforms include valve microfluidic devices^10^,^11^, microtiter plate formats such as SMART-seq2, MARS-seq, CEL- seq2 and STRT-seq^11–14^, as well as droplet microfluidics^15–17^.

An ideal platform should combine high throughput, low cost and flexibility, while maintaining the highest sensitivity and accuracy. Desirable features include imaging of each individual cell (e.g. to identify doublets and to measure fluorescent reporters), flexibility to sort cells (e.g. by FACS) and to combine multiple samples in a single run. While current valve microfluidics and microtiter plate-based formats meet most of these requirements, they are often expensive and low throughput. In contrast, droplet microfluidics achieve very high throughput and low cost per cell, but at the expense of flexibility. In particular, multistep protocols present a challenge to droplet-based systems, do not permit imaging and typically do not scale well to a large number of samples (as opposed to cells).

The adult human brain poses a particular challenge for single-cell genomics. With few exceptions, samples from human brain are only available in the form of frozen post-mortem specimens. Although good human brain banks exist, where the postmortem interval has been minimized and RNA of high quality can be extracted, it is not possible to obtain intact whole cells from such materials. Somewhat surprisingly, it has been shown that nuclei can be sufficient to derive accurate cell type information^18^, including from frozen human brain specimens^19^. However, nuclei have not yet been successfully analyzed on high-throughput platforms such as droplets or microwell arrays.

To meet these challenges, we developed an addressable nanoliter-volume microwell array platform compatible with our previously described STRT-seq chemistry, which is sufficiently sensitive to analyze both whole cells and nuclei. We designed a custom aluminum plate with outside dimensions conforming to standard microtiter plates, but with 9600 wells arranged in 96 subarrays of 100 wells each (Fig. 1a). The wells were designed with a diameter and spacing large enough to be addressable by a microsolenoid nanodispenser capable of depositing as little as 35 nL per well. With a maximum well volume of 1 μL, this facilitates efficient multi-step protocols that include separate lysis, reverse transcription and PCR steps with sufficient dilutions to avoid inhibition of later steps by the reagents used in previous steps. We modified and extensively reoptimized our 5′ STRT-seq method (Fig. S1) by introducing an additional level of indexing (‘dual index’), to allow multiplexing first within each subarray and then across the whole plate. Sequencing libraries were designed for single rather than paired-end reads, contributing to a competitive per-cell cost of the method.

**Figure 1.**
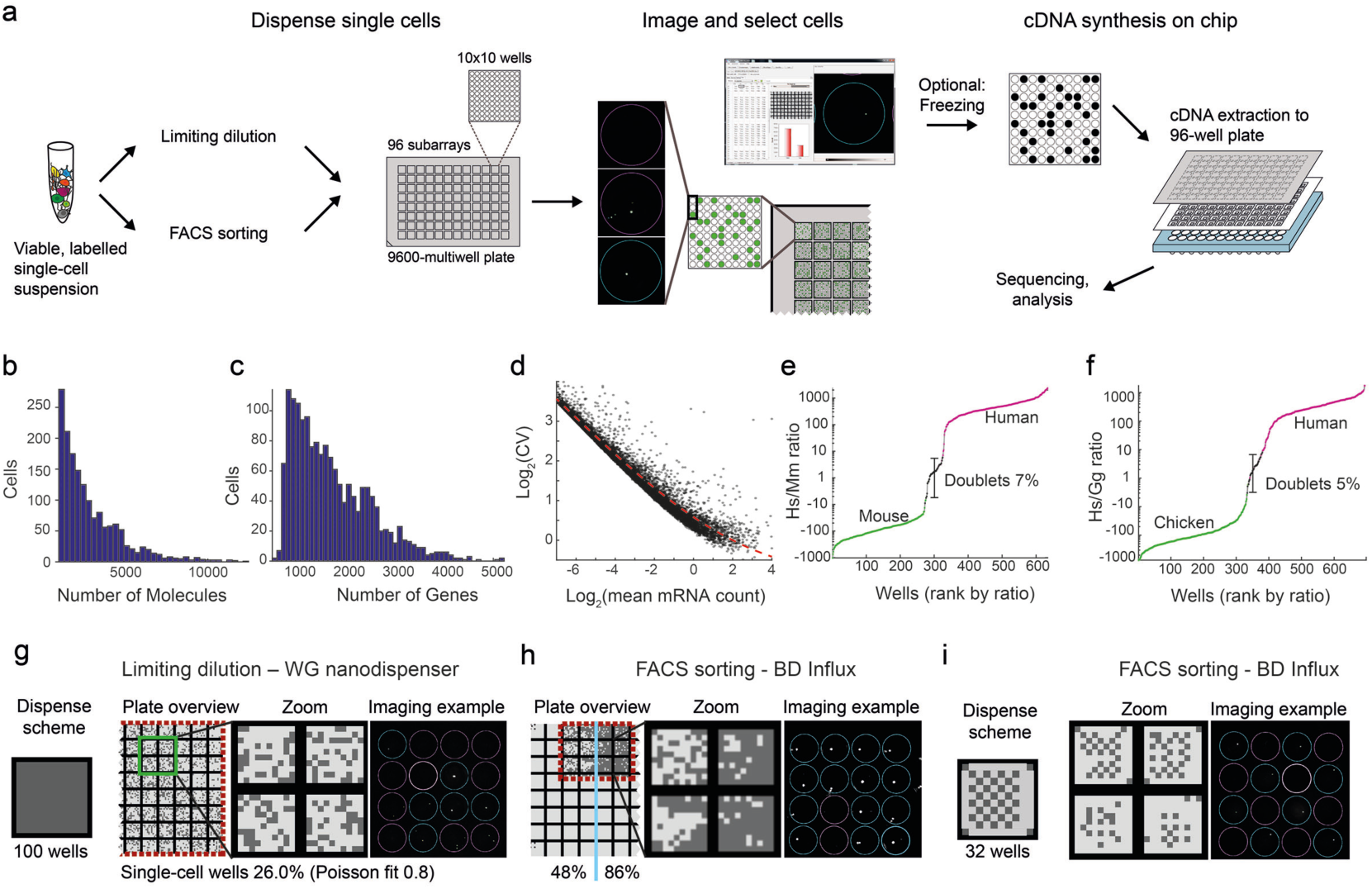
Technical performance. (a) STRT-seq-2i workflow overview. (b-c) Distribution of molecule (b) and gene counts (c) for cortex data (Fig. 2). (d) Coefficient of variation (CV) as a function of mean number of molecules *m* expressed in cortex cells. The fitted line represents an offset Poisson, *Log*_2_ *CV* = *log*_2_ (*m*^-0.5^ + 0.5) (e-f) Doublet rates as estimated by the ratio of species-specific molecules, per well, in mouse-human (e) and chicken-human (f) two-species experiments. (g-h) Single-cell well success rate when addressing 100 wells per unit by (g) limiting dilution or (h) FACS with 200 nL (left) or 50 nl PBS (right) predispensed. (i) Accuracy of FACS demonstrated by checkerboard pattern sort to 32 wells per unit.

The addressable microwell array format allows the user to process multiple samples per plate in parallel, and includes an imaging checkpoint for single-cell positive wells. Cells can be deposited by limiting dilution or by FACS, yielding up to ∼3000 or ∼7500 single cells per plate, respectively, and multiple plates can be prepared at once and frozen for later processing.

To adapt 5’ STRT-seq (also known as C1-STRT^11^) for dual indexing (STRT-seq-2i), we optimized all key steps of the protocol, including cell lysis (Fig. S2a-c), reverse transcription (Fig. S2d-e), PCR (Fig. S2f-g) and sequencing library preparation (Fig. S2h) to increase yield and quality.

In order to validate the method, we first measured its technical performance (Fig. 1b-f and Fig. S3). Using cells freshly isolated from mouse somatosensory cortex (as previously described^1^) we generated an average of 41,000 mapped mRNA reads per cell. We observed an average of 3686 detected genes, and 8706 detected mRNA molecules (Fig. S3a, distribution Fig. 1b-c) for the typical cortical pyramidal cell, comparable to previously published data^1^ with an average read depth of 500,000 mapped mRNA reads per cell (4550 genes, 17530 molecules). The number of mRNA molecules and genes detected varied greatly by cell type, indicating that this variation was dominated by biological, not technical, factors (Suppl. Fig. S3a). Noise followed an overdispersed Poisson distribution, as expected (Fig.1d).

Next, to assess possible cross-contamination between wells and subarrays, we performed mixed two-species experiments with human (Hek293) and mouse (mES) or human and chicken (DF-1) cells. Approximately 7% (mouse) or 5% (chicken) of wells contained molecules stemming from both species at roughly equal ratios, indicating true doublet wells (Fig 1e-f, Fig. S3b). These doublets were likely due to inefficient detection of poorly stained cells by imaging, since post-analysis manual inspection of putative doublet wells could not confirm the doublets. In contrast, background reads from the other species in single-cell wells was low (average 37 molecules). Therefore, ambient RNAs in the suspensions or cross-contamination occurring further downstream (e.g. during barcode indexing steps or library preparation), all contributed little to final mRNA counts.

In order to assess the performance of different cell deposition strategies, we first dispensed cells using limiting dilution, i.e. loading an average of one cell per well. We designed 32 barcodes, to allow recovery of up to 32 wells per subarray or 3072 total (slightly below the Poisson limit of 3552 cells). In practice, we observed an average single-cell fill rate of almost 2500 cells per plate (Fig. 1g, Table S1). In order to improve yield per plate, we used FACS to sort cells directly into the wells. In this mode, with optimal sorting parameters, we were able to get 86% single cells (Fig. 1h), or more than 8,000 cells per plate, although sorting that many cells was a slow process (see Methods). FACS also has other advantages, e.g. it can reduce the incidence of doublets, can be used to focus on desired rare subpopulations, and to link molecular surface properties to each individual cell by index sorting. To ensure the accuracy of FACS dispensing, we sorted cells in a checkerboard pattern, showing a deposition error rate of 4.1% of total addressed wells (Fig. 1i).

Applying the method to mouse somatosensory cortex (S1 region), in five independent experiments, we selected approximately 2200 cells (Fig. 2a). Biclustering with BackSPINv2 algorithm^9^ resolved the structure of subclasses to a similar level as reported previously (Fig 2a, Fig. S4). We detected all major cell types, including excitatory and inhibitory neurons, oligodendrocytes, astrocytes, endothelial cells, microglia and ependymal cells. We also detected known subtypes. For instance, pyramidal neurons formed distinct clusters that showed layer-specific expression profiles (Fig 2b-c)^20^. Importantly, the method showed reduced bias against cell size compared with the Fluidigm C1, demonstrated by the presence of the small oligodendrocyte precursor cells (OPC) in this dataset, which were not detected in our earlier results (Zeisel et al. 2015^1^; but see also Marques et al. 2016^9^, where OPCs were detected in a much larger dataset). Further, the full oligodendrocyte lineage was present and previously described markers (eg. *Pdgfra*, *Itpr2* and *Apod*)^9^ could be related to the maturation process from OPC to myelin-forming oligodendrocyte (MFOL) (Fig S4).

**Figure 2.**
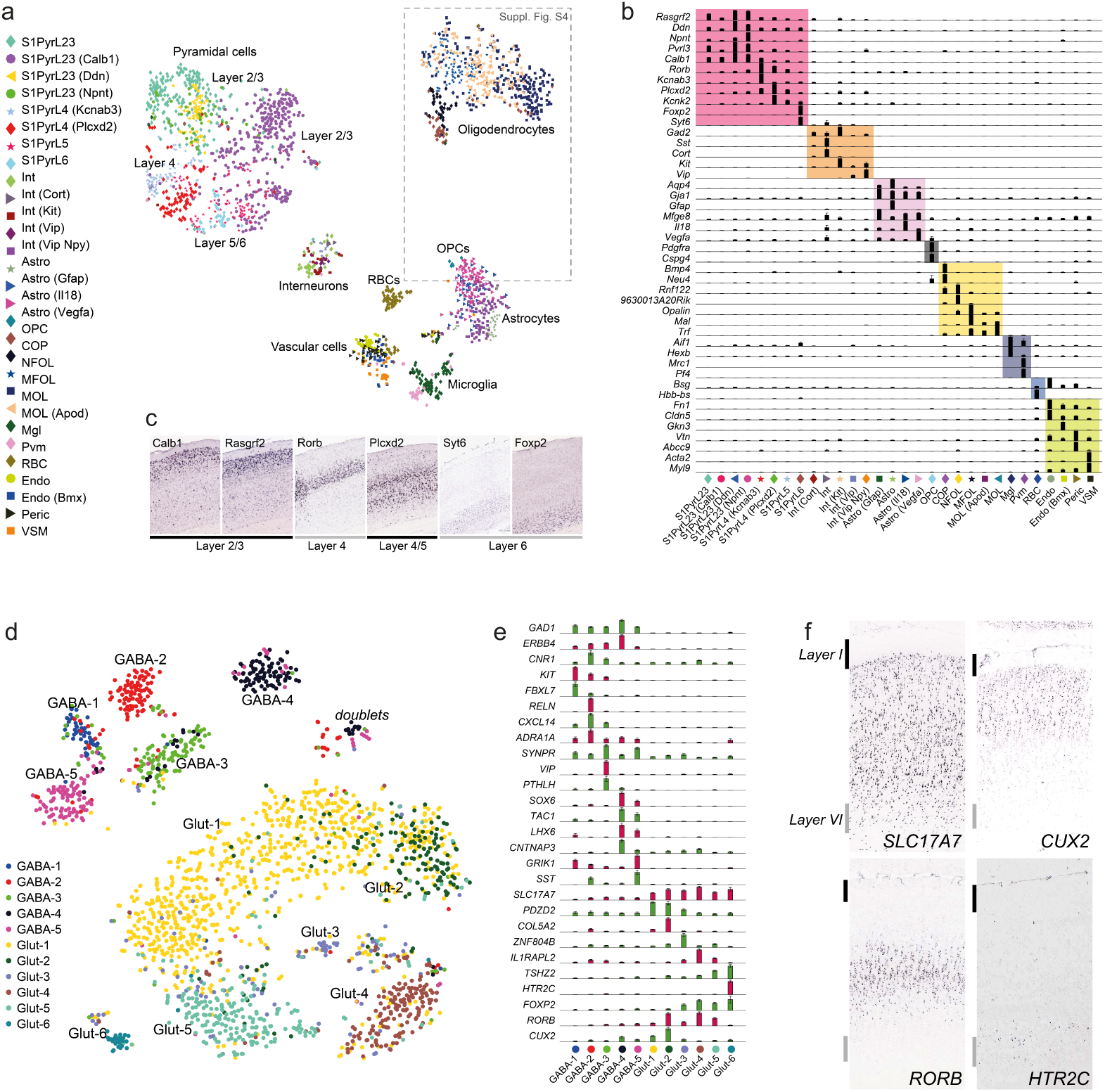
Heterogeneity of cell-types in the mouse somatosensory cortex and human temporal cortex. (a) tSNE visualization for clustering of 2192 single-cells, colored by BackSPINv2 clusters. (b) Top marker genes of each cell type presented as normalized average expression by cluster, with major cell classes overlayed by colored boxes. (c) Genes specific to pyramidal neuron subclasses by layer specificity, confirmed by *in situ* hybridization from Allen Mouse Brain Atlas. Image credit: Allen Institute. (d) tSNE visualization for clustering of 2028 post-mortem isolated neuronal nuclei from the middle temporal gyrus, colored by BackSPINv2 clusters. (e) Top marker genes of each neuronal subtype presented as normalized average expression by cluster. (f) Validation of pyramidal neuron (Glut) gene expression layer specificity, by *in situ* hybridization from Allen Human Brain Atlas. The outermost layers I and VI are indicated by strokes. Image credit: Allen Institute.

To test the versatility and sensitivity of the platform, we next used neuronal (NeuN+ FACS-sorted) nuclei isolated from a frozen post-mortem human middle temporal gyrus specimen. In a single experiment, we obtained 2028 nuclei. Despite shallow sequencing (mean <62 000 reads per cell, Fig. S5), BackSPINv2 hierarchical clustering revealed eleven distinct glutamatergic and GABAergic cell types (Fig. 2d). These were characterized by exclusive or combinatorial expression of genes (Fig. 2e), and validated by comparison with Allen Human Brain Atlas^21^ (Fig. 2f). To our knowledge, this makes STRT-seq-2i the only current high-throughput platform amenable to single-nuclei RNA-seq in human postmortem tissue.

In summary, STRT-seq-2i is a flexible and high-throughput platform for single-cell RNA-seq. It retains many of the advantages of STRT-seq, such as the use of unique molecular identifiers (UMIs) for absolute quantification, and single-read rather than paired-end sequencing for lower cost. But the transition to an addressable microwell format confers additional benefits. First, we gained the flexibility to deposit cells by dilution or by FACS, including by index sorting to track molecular surface properties of each cell and link them to the final data. We can freely deposit multiple samples (up to eight, currently) on each plate, and thereby optimize the allocation of sequencing power in complex experimental designs, something which is difficult using droplet microfluidics. Second, the plate is compatible with imaging, so that single-cell wells could be verified (albeit currently with a significant false negative rate due to imperfect cell staining), and fluorescent reporters can be linked to the final expression profile of each single cell. Third, plates can be filled, frozen – and optionally shipped – for processing at a later time (Fig. S1a-b, Alternative B). This should prove useful e.g. as single-cell RNA-seq enters clinical settings, where sample procurement and sample processing are often performed at distinct sites. Fourth, the low reaction volume and the use of single read sequencing keep costs low, and we estimate a current cost of approximately $1/cell, including 100,000 raw reads. Finally, we note that the open addressable microwell format, in contrast to droplet microfluidics, could easily be adapted to perform any multistep protocol currently implemented in regular microtiter plates (as long as they are strictly additive). This should enable a similar flexibility and throughput for other applications, such as full-length mRNA-seq (e.g. SMART-seq2^12^), whole-genome amplification, and the detection of chromatin modification^22^ and conformation^23^.

## Methods

### Cell culture

Human Hek293 and chicken DF-1 cells were cultured in complete DMEM medium. Mouse ES cells^24^ were maintained under feeder-free conditions in LIF-2i medium on 0.1% gelatin-coated culture plates^25^. The cells were trypsinized, washed, counted and assessed for cell viability.

### Animals

Male and female wild type CD-1 mice (Charles River) between postnatal days 21-37 were used. All experimental procedures followed the guidelines and recommendations of Swedish animal protection legislation and were approved by the local ethical committee for experiments on laboratory animals (Stockholms Norra Djurförsöksetiska nämnd, Sweden).

### Human post mortem tissue

Postmortem human brain tissue was provided to the Allen Institute for Brain Science by the San Diego Medical Examiner’s (SDME) office after obtaining permission for tissue collection from decedent next-of-kin. Tissue specimens were de-identified and assigned a numerical ID, and the Allen Institute for Brain Science obtained the tissue under a legal agreement that prevents SDME from sharing the key to the code or any identifying information about tissue donors. The collection and use of postmortem human brain tissue for research purposes was reviewed by the Western Institutional Review Board (WIRB). WIRB determined that, in accordance with federal regulation 45 CFR 46 and associated guidance, the use of and generation of data from de-identified specimens from deceased individuals does not constitute human subjects research requiring IRB review. All tissue collection was performed in accordance with the provisions of the Uniform Anatomical Gift Act described in Health and Safety Code §§ 7150, et seq., and other applicable state and federal laws and regulations. The tissue specimen used in this study was pre-screened for known neuropsychiatric or neuropathological history, and underwent routine serological testing and screening for RNA quality (RNA integrity number ≥7).

### Single cell suspension from mouse cortex

Single cell suspensions from adolescent mouse cortex were generated as described before^9^. Briefly, mice were anesthetized with isoflurane, perfused with ice-cold aCSF and brains collected. Brains were then sectioned using a vibratome or brain matrix and the somatosensory cortex was microdissected. Single cell suspensions were generated using the Worthington Papain dissociation system, with modifications as described^9^.

### Isolation, sorting and processing post-mortem adult human neuronal nuclei

Nuclei were isolated from a -80°C frozen tissue piece taken from the middle temporal gyrus of the cerebral cortex using Dounce homogenization, as described before^26^. Briefly, the tissue piece was thawed in homogenization buffer (10mM Tris pH 8.0, 250mM sucrose, 25mM KCl, 5mM MgCl_2_, 0.1mM DTT, 1x Protease Inhibitor (Promega), 0.4U/μl RNasin Plus RNase inhibitor (Promega) 0.1% Triton X-100) and gently homogenized with 5-10 gentle strokes using a loose pestle, followed by 5-10 strokes with a tight pestle. The homogenate was filtered through a 30μm cell strainer and nuclei pelleted by centrifugation, 10min at 900g. Nuclei were resuspended and incubated for 15min at 4°C in blocking buffer (1x PBS with 0.5% BSA and 0.2U/μl RNasin Plus RNase inhibitor), an aliquot quality assessed under the microscope, and stained with conjugated primary mouse-anti-NeuN_PE antibody, 1:500 (Millipore) rotating at 4°C for 30min. Stained suspensions were washed in blocking buffer, centrifuged 5min at 450g, transferred to FACS tubes, supplemented with 1μg/μl DAPI and sorted (FSC/SSC singlets, DAPI+, PE+). Sorted nuclei were frozen at -80°C in PBS with 10% DMSO and 0.8% BSA.

An aliquot of 100 000 NeuN+ sorted nuclei was thawed from -80°C in a 37°C water bath and quickly transferred to ice. The nuclei were diluted in 3 volumes of dilution buffer (1x PBS with 0.5% BSA and 0.5U/μl TaKaRa RNase Inhibitor) and centrifuged 5min at 1000g. Supernatant was carefully removed and the pellet was resuspended in dilution buffer.

### Cell dispense using Nanodispenser MSND

Viable single cell suspensions were stained with CellTracker Green CMFDA dye (Life Technologies) according to the manufacturer’s instructions, except incubation was 10min on ice. Suspensions were washed twice (cortex) or three times (cell lines) in respective medium and cells counted. Human nuclei were stained with Propodium Iodide ReadyProbes (Life Technologies), according to the manufacturer’s instructions. All suspensions were diluted to 20 cells or nuclei/μl in PBS (cell lines), Ca^2^+/Mg^2+^-free aCSF (mouse cortex) or dilution buffer (human nuclei). 50nl of the suspension were dispensed to all wells.

### FACS to wells using BD Influx

Before FACS, wells to be used were dispensed with 50-200nl PBS or 50nl lysis buffer and the plate was kept on ice. Cells were stained with CellTracker Green (as above) and Propodium Iodide ReadyProbes (Life Technologies), according to the manufacturer’s instructions, to discriminate dead cells. For sorting to single wells, we used BD Influx instrument (for configuration see Table S2). For a higher stability of sorting streams 140μm and 200μm nozzles were tested for efficiency. As two independent sort experiments with a 200μm nozzle setup demonstrated decreased efficiency (data not shown) the 140μm nozzle setup was used for further experiments. The gating strategy was set as follows: (1) Population of cells based on FSC-H x SSC-H profile, (2) Singlets based on FSC-H x FSC-W, (3) Singlets based on FSC-H x FSC-A. (4) One of the following options: (a) Cell-Tracker Green positive (530/40 [488nm]) or (b) Cell-Tracker Green positive (530/40[488nm]) and Propidium iodide negative (585/29[561nm]). Due to software memory limitations, only a quarter of the full layout (2400 wells) could be set at a time. Initially, two such quarter layouts, covering half the plate were created. Using the symmetric plate design, it was then turned 180° to fill the full 9600-well plate. Layouts were aligned prior to each particular experiment using a dumb plate covered with thin film, to monitor the position of 3-5 drops of Accudrop fluorescent beads. Given these limitations, we estimate a total plate fill time of below 1.5 hours.

### Imaging and Cell selection

For correct imaging positioning, fiducial fluorescent stain or highly concentrated stained cell suspension was dispensed to corner wells during cell dispense (MSND) or before FACS sort. The dispensed plate was sealed with MicroAmp Optical Adhesive Film (Applied Biosystems), centrifuged 3min at 200g and mounted upside-down on automated Nikon ECLIPSE Ti. All wells were imaged in FITC channel, using a 4x objective (4-by-4 wells per frame). Imaging took less than 15 minutes, during which the plate was cooled using ice packs, and then immediately placed on ice during image analysis (Fig. S1a Alt A), or frozen on - 80°C (Fig. S1a Alt B). Imaging files were loaded to the CellSelect Software (WaferGen) with single cell containing wells selected using varying parameters, depending on the cells used. A quick manual inspection of included and excluded wells was carried out, and if needed, analysis parameters (such as Expected Cell Size, Circularity, Brightness) were adapted. If the plate was held on ice for further processing, a maximum of 7 minutes was allowed per analysis. A final list of single cell well candidates was saved as a Filter File for dispense of all downstream reagents.

### Lysis and reverse transcription

If the plate was immediately processed (Fig. S1a Alt A), 50nl lysis mix (500nM STRT-P1- T31, 4.5nM dNTP, 2% Triton-X-100, 20mM DTT, 1.5U/μl TaKaRa RNase Inhibitor) was dispensed, followed by 3min lysis at 72°C. Then, 85nl reverse transcription (RT) mix (2.1X SuperScript II First-Strand Buffer, 12.6mM MgCl_2_, 1.79M betaine, 14.7U/μl SuperScript II, 1.58U/μl TaKaRa RNase Inhibitor, 10.5μM P1B-UMI-RNA-TSO) were dispensed and RT carried out 42°C for 90 minutes.

If the plate had been stored on dry ice or at -80°C for later processing (Fig. S1a Alt B), it first was thawed to room temperature, followed by dispense of 70nl Lysis-RT mix (1.62X SuperScript II First-Strand Buffer, 10.2mM MgCl_2_, 1.36M betaine, 425nM STRT-P1-T31, 3.4mM dNTP, 3.4% Triton-X-100, 11.9mM DTT, 8.5U/μl SuperScript II, 1.28U/μl TaKaRa RNase Inhibitor, 5.1μM P1B-UMI-RNA-TSO) and reverse transcription at 42°C for 90 minutes.

After each dispense and incubation step the plate was centrifuged for 1 minute at maximum speed (>2000g) to ensure proper collection and mixing of the reagents. For all array sealing, except during imaging, MicroSeal A film (BioRad) was used.

### Indexed PCR and extraction

After reverse transcription, 32 index primers (DI-P1A-idx[1-32]-P1B) for PCR were dispensed, such that each candidate well per 10x10 subarray received a unique index. Primer dispense was carried out in a 100nl dispense step with 2% bleach washes between each set of primers, and achieving 200nM final primer concentration in the PCR reaction. PCR mix (final concentration 1X KAPA HiFi Ready Mix supplemented with 0.2mM dNTP, 100nM DI-PCR-P1A) was dispensed in 565nl (Alt A) or 430nl (Alt B), with stock concentrations adapted accordingly. PCR was run as follows: 95°C 3min. 5 cycles: 98°C 30sec, 67°C 1min, 72°C 6min. 15 cycles: 98°C 30sec, 68°C 30sec, 72°C 6min. 72°C 5min, 10°C hold. After PCR, an extraction block was mounted on a clean 96-well plate. On top, the plate was mounted, upside down, to align with the extraction block. The assembly was centrifuged 5min at maximum speed (>3000g). The 96-well plate containing pooled, index amplified cDNA was assessed for quality on Bioanalyzer, sealed and stored at -20°C.

### Tagmentation and isolation of 5’ fragments

Amplified cDNA was tagmented using 96 transposomes, each with a different index to target cDNA from one subarray each. Transposome stocks were assembled (6.25μM barcoded adapter (STRT-Tn5-Idx[1-96]), 6.25μM Tn5 transposase, 40% glycerol), 37°C 1h, and concentration adapted according to Tn5 activity, if needed. Tn5 reactions were assembled with 3μl transposome and 2μl amplified cDNA, in a total 20μl 1x CutSmart buffer (NEB), and incubated at 55°C 20min. 100μl Dynabeads MyOne Streptavidin C1, washed according to the manufacturer’s instructions, were diluted 1:20 in BB buffer (10mM Tris HCl pH 7.5, 5mM EDTA, 250mM NaCl, 0.5% SDS), added to the tagmentation reaction 1:1, and incubated at room temperature 15min. After incubation, all samples were pooled to one tube, washed twice in TNT (20mM Tris HCl pH 7.5, 50mM NaCl, 0.02% Tween-20) and resuspended in 50μl TNT. Remaining adapters were cleaned by adding 10μl ExoSAP IT (Affymetrix) and incubating 15min 37°C, followed by two more washes in TNT and one careful wash in EB. The single-stranded library was eluted in 50μl nuclease-free water by incubating 10min at 70°C and collecting the supernatant to a new tube. The library was then bound to 1.5X volumes AMPure beads (Beckman), supernatant removed, the beads resuspended in Second PCR mix (1X KAPA HiFi Ready Mix supplemented with 200mM 4K-P1_2ndPCR, 200nM P2_4K_2ndPCR), and cycled (95°C 2min. 8 cycles: 98°C 30sec, 65°C 10sec, 72°C 20sec. 72°C 5min, 10°C hold). The supernatant was transferred, cleaned with 0.7X volumes of AmPure beads and eluted. The eluate was bound to 0.5X volumes of AmPure beads, the supernatant transferred, again bound in 1X volume of AmPure beads, and eventually eluted in EB.

### Illumina high-throughput sequencing

The quality and molar concentration of libraries were assessed by Bioanalyzer. Sequencing was performed on Illumina HiSeq2000 or 2500 with Single-End 50 cycle kit using the Read1 DI-Read1-Seq, Index 1 STRT-Tn5-U and Index 2 DI-idxP1A-Seq. Reads of 45 bp were generated, expected to start with a 6 bp UMI, followed by 3 guanidines, and the 5’ transcript. The two index reads of 8 and 5 bp corresponded to Index 1 (subarray barcode) and Index 2 (well barcode), respectively.

### Data analysis

Reads flagged as invalid by the Illumina HiSeq control software were discarded. In the remaining reads, any 3′ bases with a quality score of B were removed. If any of the six UMI bases in the 5' end had a Phred score <17 the read was discarded, else the UMI was cut away and saved. The following three bases had to be G for the read to be kept. These, as well as any additional up to totally nine (presumably template-switch derived) G:s were cut away. Reads were discarded if the remaining transcript-derived sequence ended in a poly(A) sequence leaving <25 bases, or if the remaining sequence consisted of fewer than six non-A bases or a dinucleotide repeat with fewer than six other bases at either end. The reads were sorted by the well, as defined by the combined two index read barcodes.

Alignment was performed using the Bowtie aligner^27^, allowing for up to three mismatches and up to 24 alternative mappings. Reads with no alignments were realigned against an artificial chromosome, containing all possible splice junctions arising from the exons defined by the UCSC transcript models^28^. The coordinates of aligned splice junctions were translated back to the corresponding genomic positions.

For expression level calculation, the exons of each locus with several transcript variants were merged to a combined model representing total expression of the locus. To account for incomplete cap site knowledge, the 5′ ends of all models were extended by 100 bases, but not beyond the 3′ end of any upstream nearby exon of another gene of the same orientation.

The annotation step was performed separately for each well. For each genomic position and strand combination the number of reads in each UMI was counted. Any multiread that mapped to some repeat outside exons was assigned randomly as one of these repeats and did not contribute to the transcript corresponding to the exon. Else, if the multiread mapped to some exon, and not to any repeat outside exons, it was assigned to the exon where it was closest to the transcript model 5′ end. If it had no exon mapping, it was assigned randomly at one of the mappings. The total number of molecules at each mapping position was determined by the number of distinct UMIs observed. Any UMI represented by only a single read was excluded, in order to reduce false molecules due to PCR and sequencing errors. The raw UMI count was corrected for the UMI collision probability as described^29^.

The human nuclei samples were analyzed similarly, but all reads mapping anywhere within the whole locus, defined as the region (including actual introns) from the start of the 100 base 5'end extension of the first exon up to the end of the last exon, were counted as exon-derived.

### Clustering and analysis mouse cortex cells

Analysis of mouse cortex samples included the following steps: (1) Loaded all 7769 cells. (2) Filtered on 800-2000 total molecules per cell and ratio total molecules/total genes >1.2, resulting in 6449 cells. (3) Excluded doublets identified by co-expression of known marker genes (*Stmn2* – neurons, *Mog* – oligodenderocytes, *Aqp4* – astrocytes, *Fn1* – endothelial, *C1qc* – microglia), 5514 cells retained. (4) Permuted order of cells and genes. (5) Clustering by BackSPINv2 with following parameters: splitlev = 7; Nfeture = 300; Nfeture1 = 500; N_to_backspin = 10; N_to_cells = 500; mean_tresh = 0.01; fdr_th = 0.3; min_gr_cells = 5; min_gr_genes = 10; stop_th = [0.5,0.5]; flag_val_stip = 1;. (6) Manual inspection of clustering results and separation of cells to neurons and non-neurons. Clusters that showed no specific marker, interpreted as low quality, and cells originating from a Hek293 sample (Table S1) were removed (1882 cells removed, Fig S4a) (7) Separate clustering of neurons and non-neurons with following parameters: splitlev = 6; Nfeture = 200; Nfeture1 = 800; N_to_backspin = 10; N_to_cells = 500; mean_tresh = 0.01; fdr_th = 0.3; min_gr_cells = 5; min_gr_genes = 20; stop_th = [0.5,0.5]; flag_val_stip = 1;. (a) For neurons, this resulted in 35 clusters, we excluded 12 clusters for which no specific marker was obtained and merged some of the remaining 23 clusters into final 13 clusters (Fig S4b). (b) For only non-neurons the same BackSPINv2 parameters resulted in 33 clusters, we excluded 7 clusters without specific marker and merged some of the remaining 26 clusters into 17 final clusters (Fig S5). tSNE projection^30^ was used for visualization only. We used the following parameters: 410 genes, perplexity 20, PCA components 20, epsilon 100, distance correlation number of iterations 1000 (Matlab code https://lvdmaaten.github.io/tsne/). In the heatmap (Fig S4) we used the same 410 genes. Color on heatmaps represent normalized expression (log-transform per gene: mean=0, standard deviation=1) and are saturated 1-99%.

### Clustering and analysis human cortex nuclei

Analysis of human cortex nuclei included the following steps: (1) Loaded all 2842 cells. (2) Filtered out cells with less than 500 detected molecules (2306 cells retained). (3) Removed genes expressed in less than 20 cells, or more than 60% of cells. (4) Normalized each cell to total 3000 molecules. (5) Clustering by BackSPINv2 using the following parameters: splitlev = 6;Nfeture = 500;Nfeture1 = 500;N_to_backspin = 10;N_to_cells = 800; mean_tresh = 0.01; fdr_th = 0.3;min_gr_cells = 5;min_gr_genes = 5;stop_th = [0.5,0.5];flag_val_stip = 2. Clustering resulted in 31 clusters. (6) One cluster expressing glial genes was removed, the rest were manually merged into 11 clusters. For visualization, we selected top enriched genes, as described above. tSNE visualization was run as above, with the following parameters: initial PCA dimension = 20, perplexity = 2, epsilon = 100, correlation as distance, maximum iterations = 1000.

### Two-species analysis

A combined two-species bowtie-1^27^ alignment index was constructed from transcript models as defined by the UCSC refFlat table data^31^. In order to obtain equally-sized and similar representations of the two transcriptomes to compare, we naively restricted the analysis to the transcripts that had identical names in the two species (disregarding upper/lower case). Illumina HiSeq reads were processed as follows: Reads ending in a poly(A) sequence leaving less than 25 alignable 5' bases were discarded. Any 3’ bases with a quality score of B were removed. If the remaining sequence consisted of fewer than six non-A bases or a dinucleotide repeat with fewer than six other bases at either end, the read was discarded. The filtered reads were aligned to the bowtie-1 index allowing for up to 3 mismatches. A read was considered unequivocally belonging to a species and counted if it had a perfect match to a transcript of that species, but no match in the other species, also when allowing up to 3 mismatches. Molecule counts were obtained as the number of distinct six nucleotide long unique molecular identifiers (UMIs) of the reads aligned at each position. UMIs represented by a single read were not counted, since our previous experiments have shown that these to a large extent are artifacts stemming from PCR and sequencing errors and result in overestimates of molecule counts.

### Primer sequences

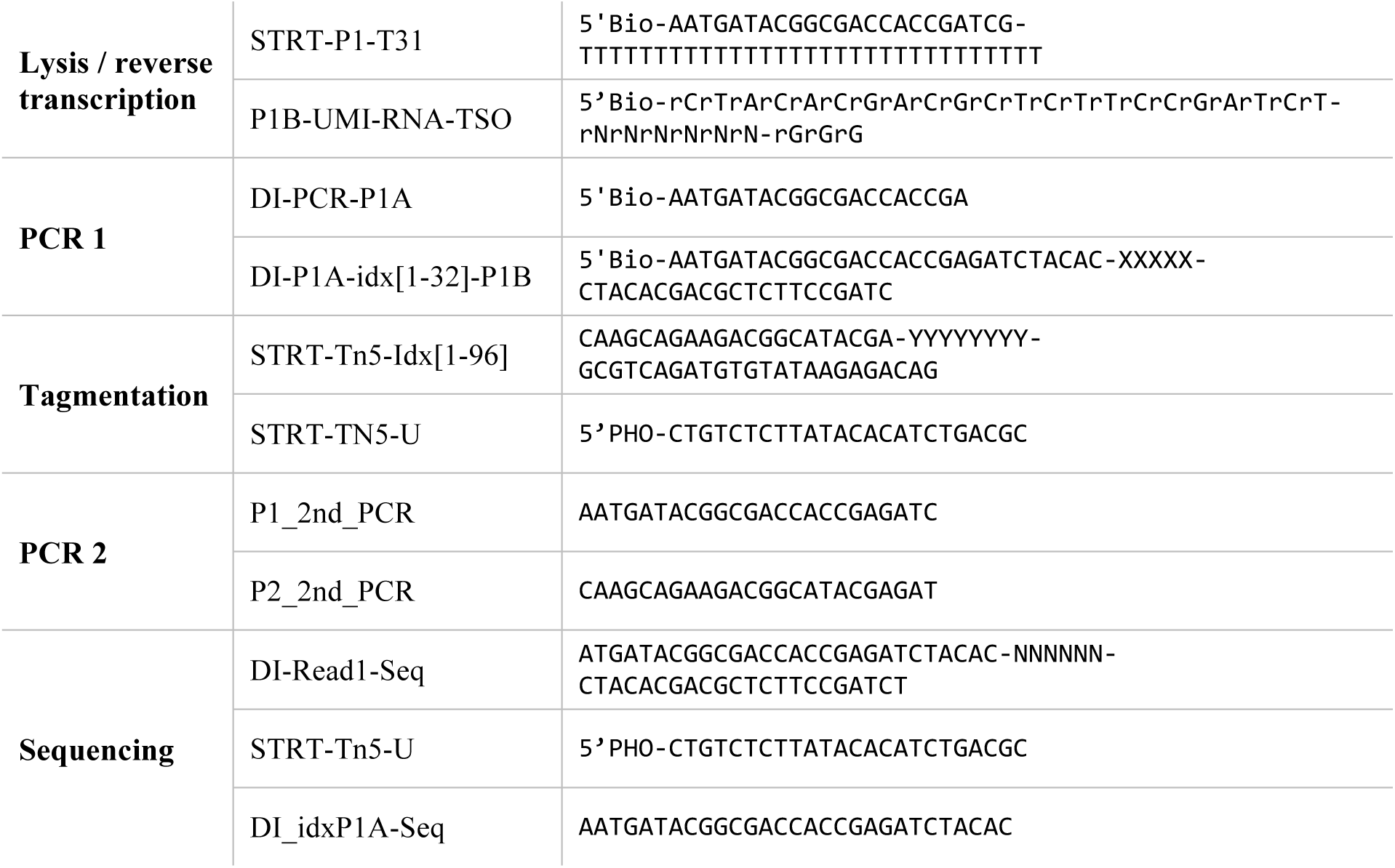

## Author contributions

A.Z, M.S, S.L, H.Ho, H.Hu, G.L.M and J.D conceived and designed the method. H.Hu, P.Li, J.D, S.H, M.S and A.Z designed and engineered the microarray. H.Ho, R.H, J.M, A.H and A.Z performed experiments. M.R, R.H and E.L provided materials. H.Ho, H.Hu, P.Lö, A.Z and S.L analyzed data. H.Ho, A.Z and S.L drafted the manuscript, with input from all authors.

## Acknowledgements

We thank Feng Zhang for mouse ES cells. We thank Anna Johnsson for lab management and Anna Juréus for technical assistance and sequencing.

